# Oligomerisation of THAP9 transposase: role of DNA and amino-terminal domains

**DOI:** 10.1101/2020.10.06.328815

**Authors:** Hiral M. Sanghavi, Sharmistha Majumdar

**Author notes:** Correspondence should be addressed to Sharmistha Majumdar.

## Abstract

Active DNA transposases like the Drosophila P element transposase (DmTNP) undergo oligomerisation as a prerequisite for transposition. Human THAP9 (hTHAP9) is a catalytically active but functionally uncharacterised homologue of DmTNP. Here we report (using co-IP, pull down, co-localization, PLA) that both the full length as well as truncated hTHAP9 and DmTNP (corresponding to amino-terminal DNA binding and Leucine-rich coiled coil domains) undergo homo-oligomerisation, predominantly in the nuclei of HEK293T cells. Interestingly, the oligomerisation is shown to be partially mediated by DNA. However, mutating the leucines (either individually or together) or deleting the predicted coiled coil region did not significantly affect oligomerisation. Thus, we highlight the importance of DNA as well as the amino-terminal regions of both hTHAP9 and DmTNP, for their ability to form higher order oligomeric states. We also report that Hcf-1, THAP1, THAP10 and THAP11 are possible protein interaction partners of hTHAP9. These studies lead to several questions about the different putative oligomeric states of hTHAP9 and how they may be related to its yet unknown physiological role as well as interaction partners.

## Introduction

Transposable elements (TEs) are DNA segments that can change their position in the genome irrespective of any sequence homology between the excision and integration sites. This movement is aided by proteins known as transposases, often encoded by TEs. Numerous genome sequencing projects unanimously provide evidence that TEs, which are present in all kingdoms of life, form a substantial portion of the genome of a species (1) and may provide raw material for generation of new protein coding genes in the host genome by the process of evolutionary tinkering or exaptation (2). Some of the most widely known examples of such TE-derived domesticated or neo-functionalized genes are Rag1, Zbed1, SETMAR (3).

Human THAP9 (hTHAP9), a recently discovered homolog of the well characterised Drosophila P element transposase (DmTNP), may be domesticated or neo-functionalized. The hTHAP9 protein, of unknown cellular function, is a member of the twelve member human THAP family and is 25% identical and 40% similar throughout the length of DmTNP (4). DmTNP, an 87kDa active DNA transposase (5), is a member of subclass I of DNA transposons which have DNA signatures like Terminal Inverted Repeats (TIRs), Sub-Terminal Inverted Repeats (sTIRs), Target Site Duplication (TSDs) as well as the gene encoding the transposase (if autonomous). If a transposon is active, the corresponding genome may have many copies of the inserted transposon that may be full length, truncated or internally deleted (6). Interestingly, hTHAP9 was found to retain catalytic activity despite being a single copy gene with no TIRs flanking 1 kb of the hTHAP9 gene (chromosome 4) (4).

DNA transposons, which are flanked by specific DNA sites (TIRs) on either transposon end, are mobilised by transposase proteins that bind and cleave these TIRs. Many DNA transposases (both prokaryotic or eukaryotic) need to multimerize to form an active transpososome (7, 8) such that two different subunits interact with either transposon end and loop out the DNA in a synaptic nucleoprotein complex. However, the mode of oligomerisation is transposase-specific and the oligomerisation domain is formed by different structural cores. For example, an alpha helical domain in Hermes transposase (9), a leucine zipper motif in *IS911* transposase (10) or a coiled coil motif in IS2 transposase (11). Some transposases are inherently monomeric (Tn5, MuA) but oligomerise upon binding target DNA (Suppl. Fig. 1b) (8). On the other hand, the Mos1 and *Hermes* transposases are inherently dimeric or octameric respectively (Suppl. Fig. 1a). Domesticated transposases like PiggyMac (12) or TE derived proteins like RAG1 and RAG2 are also known to form functional oligomers (Suppl. Fig. 1c) (13, 14). Thus, different transposases adopt distinct strategies to assemble catalytically active nucleoprotein complexes.

DmTNP has been reported to exist as a tetramer in the early stages of transposition: as a pre-formed tetramer before binding DNA or GTP and when it forms a single end complex (SEC) or synaptic paired end complex (PEC) after binding one or both transposon DNA ends respectively (15). GTP promotes synapsis i.e formation of PEC (16) and has been shown to directly interact with and possibly position transposon DNA for catalysis (17). The single or double end cleaved donor complex (CDC) of DmTNP formed after cleavage of transposon DNA, are also tetrameric (15). On the other hand, the recent cryo-EM structure of the post transposition strand transfer complex (STC) comprises of a DmTNP dimer bound to GTP, pre-cleaved donor and target DNA (Suppl. Fig. 1d) (17). It has been speculated that the tetrameric DmTNP PEC/CDC loses two subunits after the excision of P element donor DNA, to form the dimeric STC that carries out transposon integration in a target site (17).

In addition to being indispensable for the DNA transposition process, oligomerisation of the transposase protein has been shown to be an important mode of regulation of transposition. For example, the internally deleted KP element, encodes a 24kD Type II repressor protein known as KP repressor (18) which shares 100% sequence identity with DmTNP for the first 199 residues and inhibits P element transposition at multiple levels: it binds to multiple DmTNP binding sites near the P-element termini, thus making these sites unavailable for DmTNP binding (19); it may also form a heteromultimeric complex with DmTNP, thus making DmTNP less available for transposition (20). The IS50 element also encodes a transposition inhibitor that lacks the amino terminal 55 amino acids and DNA binding domain (DBD) but interacts with the full length transposase via oligomerisation (21). Interestingly, formation of higher order oligomers of Mos1 (22) and Hsmar1(23) have been suggested to form aggregates which lead to inactivation of transposase (Mos1) or decrease the catalytic activity of the transposase (Hsmar1).

In this study, we use a combination of *in silico* approaches to predict the oligomerisation regions in both DmTNP and hTHAP9. Biochemical analysis establish that the full length hTHAP9 protein can oligomerise like DmTNP. Interestingly, truncated versions of both DmTNP and hTHAP9, which included the DBD and the predicted coiled coil region, retained the ability to oligomerise. However, deletion of the predicted coiled coil region does not disrupt oligomerisation. Finally, DNaseI cleavage assays indicate that the oligomerisation of both DmTNP and hTHAP9 may be partially mediated by DNA.

The physiological functions as well as the cellular interaction partners of hTHAP9 are unknown. Given hTHAP9’s ability to oligomerise, we decided to explore its possible interacting proteins by using a candidate based approach. THAP1, THAP10 and THAP11 from the human THAP family, were chosen as candidates based on predictions by the STRING database (24). All THAP proteins except THAP0, THAP4 and THAP8, have the predicted Hcf-1 binding consensus motif (HBM): ((D/E)HXY) (25). Hcf-1 is a key regulatory protein involved in diverse cellular pathways like cell cycle progression, embryonic stem cell pluripotency and stress response (26). We demonstrate that hTHAP9 colocalises with human THAP1, THAP10, THAP11 and Hcf-1.

Overall, we demonstrate that hTHAP9, like its homolog DmTNP, can undergo homo-oligomerisation, which is partially mediated by DNA. Moreover, hTHAP9 can also hetero-oligomerise with several members of the human THAP family as well as Hcf1.

## Methods

### *In silico* prediction of coiled coil domains

The coiled coil domains were predicted as described earlier (27). Briefly, the amino acid sequence of full length hTHAP9 and DmTNP were submitted to freely available softwares. In a given amino acid sequence, COILS predicts the region with some probability of forming coiled coils (28). Three different scanning windows (of length 14, 21 or 28 amino acids) are considered. The predictions using a 28 residue window are preferred for a less studied protein sequence. The predictions with a probability of more than 50% (0.5) are considered more promising. JPRED (29) was used to predict the secondary structures of the predicted coiled coil regions in hTHAP9 and DmTNP. The secondary structure predictions were correlated with the protein structure predicted by I TASSER (30). Further, these were used to predict interactions amongst themselves using DRAW COIL (31).

### Plasmid constructs

The full length hTHAP9 cDNA was cloned in frame with carboxy-terminal 1XFLAG tag (THAP9-F) or 1XHA (THAP9-H) tag by Genei Laboratories Private Ltd.

The hTHAP9 open reading frames encoding amino acids 1-176 (H1) and 1-165 (H2) were amplified from full length hTHAP9 cDNA. Carboxy-terminal 3XHA tags were added using two overlapping PCR reactions and the inserts were subcloned into pcDNA3.1+ using NotI and XbaI restriction enzymes (H1-H, H2-H). H1 (F-H1) and H2 (F-H2) were also cloned with amino-terminal 3XFLAG tags into pcDNA3.1+ using the same method.

The DmTNP open reading frame encoding amino acids 1-140 (D) was amplified from full length DmTNP cDNA with carboxy-terminal 3XHA tag (D-H) or amino-terminal 3XFLAG tag (F-D) using the same method as that for hTHAP9.

Vectors containing full length human Hcf-1 cDNA with amino-terminal HA tag (Plasmid #53309; H-HCF-1) and full length human THAP11 cDNA with carboxy-terminal FLAG tag (Plasmid #28020; F-THAP11) were obtained from Addgene. Vectors containing full length human THAP1 cDNA with carboxy-terminal 1X FLAG tag (THAP1-F) and full length human THAP10 with amino-terminal 3XFLAG tag (F-THAP10) were generous gifts from Professor Winship Herr’s lab.

### Antibodies

Antibodies used in this study are as follows: For IPs, pull downs and Western blots, anti-HA (Invitrogen, 71-5500) and anti-FLAG (Sigma, F7425) antibodies. For Western blotting, anti-rabbit (GE, NA934V) and anti-mouse (GE, NA931V) HRP-conjugated secondary antibodies. For immunofluorescence, anti-HA (Sigma, SAB1306169) and anti-FLAG (Sigma, F3165) primary antibodies and anti-rabbit (Thermofisher, A11037) and anti-mouse (Thermofisher, A11029) Alexa-fluor secondary antibodies.

### Cell culture and transfection

HEK293T cells were grown in DMEM (Himedia, AL007A), supplemented with 10% FBS and L-Glutamax (Invitrogen) at 37°C and 5% CO2. To check for protein expression using immunoblotting assay, 10^6 cells were seeded per well of a six-well plate and transfected, using Lipofectamine 2000 (Invitrogen, 11668019) according to manufacturer’s instructions, with 2.5 ug each of different constructs (THAP9-H, F-H1, H1-H, F-H2, H2-H, F-D and D-H) and 5ug of THAP9-F. For co-immunoprecipitation, 10^6 cells were co-transfected with 2.5 ug of each pair of plasmids (F-H1 and H1-H), (F-H2 and H2-H), (F-D and D-H) separately and the co-transfected cells were harvested 48 h post-transfection. For pull down assays, 10^7 cells were independently transfected with 2.5ug of THAP9-H (24h) and 5ug of THAP9-F (48h).

### Immunofluorescence

About 200 HEK293T cells were seeded on coverslips. F-D and D-H, F-H1 and H1-H and F-H2 and H2-H were separately co-transfected when the cells were spatially isolated from each other. The co-transfected cells were washed twice with 1X PBS 48h post transfection, following which they were fixed on coverslips using 4% formaldehyde in PBS for 10 min at 37°C. Excess formaldehyde was washed off with 1X PBS. The fixed cells were then permeabilized using a permeabilization buffer (0.1% Triton-X-100 in PBS) for 10 min at 37°C, blocked using blocking buffer (10% serum and 0.05% Triton-X-100 in PBS) for 1h at 37°C in a humid chamber, followed by incubation with 1: 100 dilution of primary antibodies (Rabbit Anti-HA, SAB1306169 and Mouse Anti-FLAG, F3165) for 2h at 37°C in a humid chamber and finally 1: 1000 dilution of mouse Alexa Fluor 594 and rabbit Alexa fluor 488 secondary antibodies for 2h at room temperature in the dark. After washing off excess secondary antibodies, the cells were mounted in a DAPI containing mounting media (Ab104135) and stored at 4°C until imaging. Samples were examined using a Leica confocal microscope (10× eyepiece and 40× objective). As the HEK293T cells tend to be in different planes of focus in each plane of view, many serial Z stack images of the individual plane of view were recorded. These images were flattened to a 2D image using the Z project function of FIJI software (32) which was further used to generate the images represented in this study.

The cells co-transfected with THAP9-H and (i) THAP1-F (ii) F-THAP10 or (iii) THAP11-F were fixed with formaldehyde 24h post transfection because THAP9-H was found to express the best after 24h.

Pearson’s R value, calculated for each of ten chosen cells in the merged panel per sample, was compared to that of the control (lipofectamine treated cells) using unpaired T test with Welch correction, in Prism 7.0c (Graphpad).

### Pull down assays

The cells transfected with THAP9-H and THAP9-F were harvested separately using chilled 1X PBS. The cell pellet was resuspended in an equal amount of RIPA lysis buffer. The THAP9-H cell lysate was incubated with 5ug of HA antibody (Invitrogen, 71-5500) at 4°C for 14-16h on a rotating tumbler followed by incubation with protein G sepharose beads (GE, 17061801) for 2h at 4°C. The G sepharose beads bound to THAP9-H protein were separated by centrifugation at 1000 rpm for 2 min, washed twice with 1X PBS followed by incubation with THAP9-F lysate for 6-8h at 4°C on a rotating tumbler. The G sepharose beads bound to protein were separated by centrifugation at 1000 rpm for 2 min and washed twice with 1X PBS. The protein bound to beads was eluted in 5X SDS gel loading buffer (250mM Tris Cl pH 6.8, 10% SDS, 30% glycerol, 0.5 M DTT, 0.02% Bromophenol blue) by boiling at 95°C for 10 min. The beads were separated from the bound protein by spinning down the beads in gel loading buffer at 13000 rpm 10 min.The supernatant was collected in a fresh tube and run as the bound fraction on a 10% SDS polyacrylamide gel.

In a separate experiment, the THAP9-F cell lysate was incubated with 5ug of FLAG antibody (Sigma, F7425) followed by incubation with THAP9-H lysate and processed as mentioned above.

### Co-Immunoprecipitation

The co-transfected cells were harvested using chilled 1X PBS. The cell pellet was resuspended in an equal amount of RIPA lysis buffer. The whole cell extract was pre-cleared by incubating with 5ug rabbit IgG (Novus Biologicals, AB-105-C) for 1h on ice followed by 75-100 ul protein G sepharose beads for 1h at 4°C on a rotating tumbler. The pre-cleared lysate thus obtained was divided into 4 fractions: 20% was used as gel input, while 25% fractions were incubated with anti-FLAG (Sigma, F7425), anti-HA (Invitrogen, 71-5500) or IgG antibodies for 16-18h at 4°C, followed by incubation with G sepharose beads for 2h at 4°C. The protein G sepharose beads bound to protein were separated by centrifugation at 1000 rpm for 2 min. The supernatant was collected in a separate tube and run as the unbound fraction on the gel. The beads were washed thrice with 1X PBS. The protein bound to beads was eluted as described above for the pull-down experiments and run on a 19% SDS polyacrylamide gel.

### Immunoblot analysis

Immunoprecipitated samples were separated by 19% SDS-PAGE, transferred to PVDF membrane and blocked with 5% serum in TBST for 1h at room temperature. Membranes were incubated overnight at 4°C with 1:2000 dilution of primary antibodies (FLAG; Sigma, F7425 and HA; Invitrogen, 71-5500) followed by incubation with 1:5000 dilution of HRP-coupled secondary antibodies for 2h at room temperature and detected using enhanced chemiluminescence (Pierce, PI32106).

### Co-Immunoprecipitation followed by DNaseI digestion assay

The protein bound beads obtained after co-immunoprecipitation were washed thrice with 1X PBS. These beads were then resuspended in equal amounts of PBS+ DNaseI digestion buffer with 3 ul (6U) of DNAseI (M0303S, NEB) and incubated at 37°C for 40min - 1h and separated by centrifuging at 2000 rpm for 2 min. The samples were then processed like the co-immunoprecipitated samples above and run on 19% SDS-PAGE followed by western blotting.

### Proximity Ligation Assay (PLA)

200 HEK293T cells were seeded on coverslips and co-transfected with F-D and D-H, F-H1 and H1-H or F-H2 and H2-H when the cells were spatially isolated from each other. As a control, HEK293T cells were treated with only lipofectamine. The co-transfected cells were washed twice with 1X PBS 48h post transfection, following which they were fixed on coverslips using 4% formaldehyde in PBS for 10 min at 37°C. Excess formaldehyde was washed off with 1X PBS. The fixed cells were then permeabilized using 0.1% Triton-X-100 in PBS for 10 min at 37°C. After the removal of excess Triton-X-100, PLA (using Duolink^®^ in situ Red Starter Kit; Sigma Aldrich DUO92101) was performed by blocking the permeabilized cells using DuoLink blocking solution in a humidity chamber at 37°C for 1h. The excess blocking solution was removed by tapping the coverslips. Primary antibodies (Rabbit Anti-HA Ab, SAB1306169 and Mouse Anti-FLAG Ab, F3165) were diluted 100 fold in the antibody diluent solution and incubated for 2h in the humidity chamber at room temperature. Excess primary antibody was washed twice with 1X wash buffer A. The PLA probes diluted in antibody diluent solution were then incubated with cells for 1h at 37°C. The excess PLA probes were removed by washing twice with 1X wash buffer A, followed by incubation with ligase at 37°C for 30 min. The signal obtained after ligation is intensified by adding the polymerase following the removal of excess ligase using 1X wash buffer A and incubating at 37°C for 100 min. Excess polymerase is removed by washing twice with 1X wash buffer B and 0.01X wash buffer B. The excess buffer is removed from the coverslips, which are then mounted on slides using mounting media (Ab104135). Samples were examined using a Leica confocal microscope (10× eyepiece and 40× objective). The PLA foci (red) were counted for D, H2 and control (in 100 cells) and for H1 (in 70 cells).

### Site Directed Mutagenesis

The Q5 Site directed mutagenesis kit (NEB, E0554S) was used to make the following mutations All mutant constructs were verified by Sanger sequencing.

### Statistical significance

Statistical significance was calculated using Prism 7.0c (Graphpad). Briefly, unpaired student’s t test with Welch’s correction was used to calculate statistical significance of (i) co-localization of two proteins in Fig. 1,4,9 (ii) in PLA, occurrence of interactions upon over-expression of respective constructs in comparison to the no-overexpression control. The paired student’s t test was used to calculate statistical significance between the ratio of nuclear and extranuclear interactions because of the over-expression of respective constructs in PLA. The bar graphs have been displayed as Mean ± SD.

**Figure 1.**
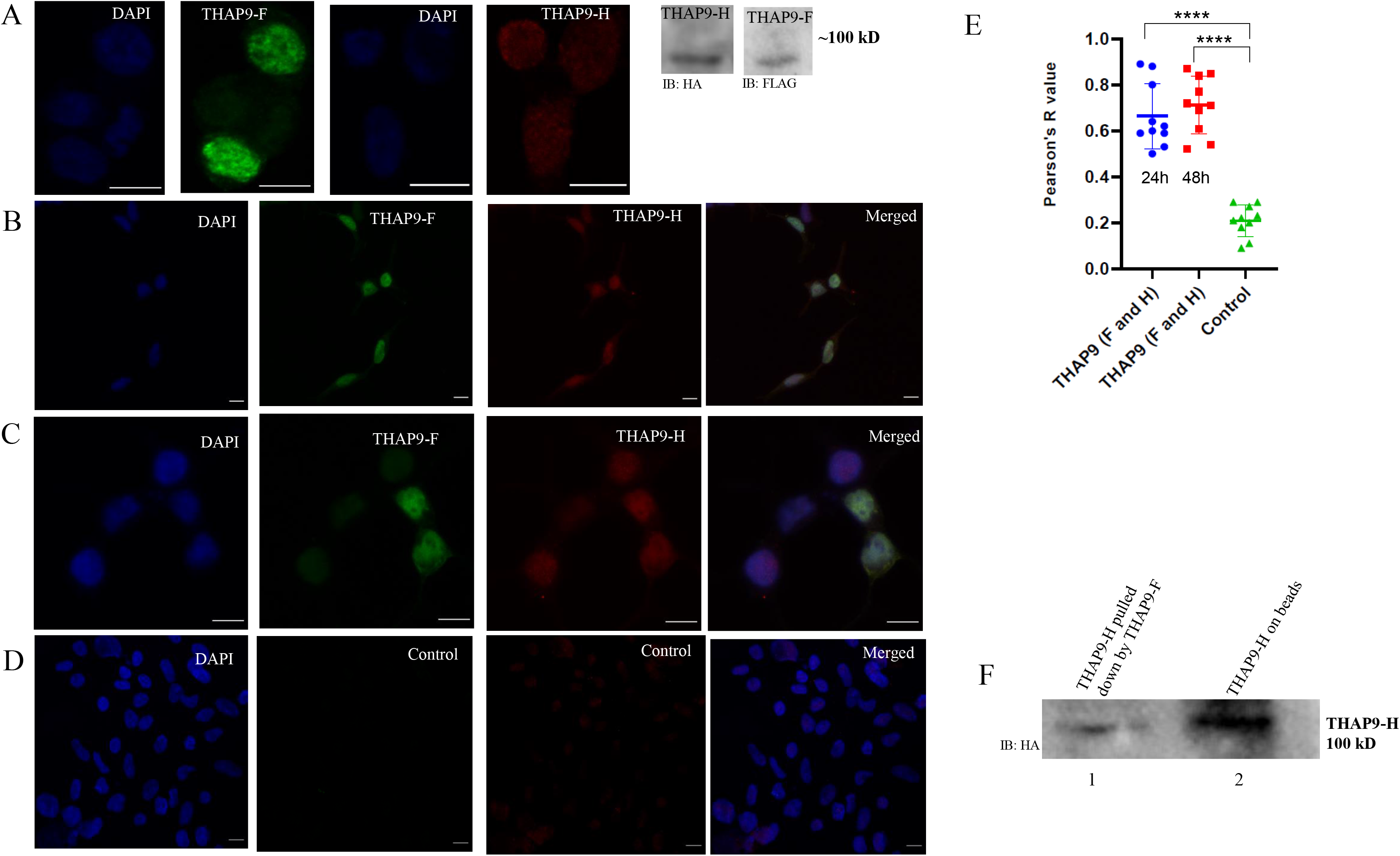
Human THAP9 homodimerises in the nuclei of HEK293T cells. (A) HEK293T cells transfected individually with THAP9-F or THAP9-H were immunostained (1st 4 panels) for FLAG (green; 1st 2 panels) or HA (red; 3rd and 4th panels) and nuclei were counterstained with DAPI (blue). Immunoblotting (5th panel; anti-HA, anti-FLAG) of corresponding whole cell extracts harvested 24 h (THAP9-H) and 48h (THAP9-F) after transfection HEK293T cells co-transfected with THAP9-H (red) and THAP9-F (green) were immunostained and observed (B) 24h post co-transfection (C) 48h post co-transfection (D) control HEK293T cells treated with lipofectamine alone for 48h. The images shown here were captured by Leica confocal microscope (10× eyepiece and 40× objective), scale bars= 10 microns. (E) Pearson’s R-value (no threshold) for co-localization of THAP9-H and THAP9-F corresponding to 1B or 1C when compared to 1D. The number of cells counted for each sample n=10. (F) Immunoblotting (anti-HA) of pull down of THAP9-H with THAP9-F (lane 1) and THAP9-H bound to beads (lane 2).

## Results

### Human THAP9 forms a homodimer

DmTNP has been reported to oligomerise as a prerequisite for active P element transposition (15). However, it is not known if hTHAP9, the less studied homolog of DmTNP, forms oligomers. Thus, the possibility of hTHAP9 forming oligomers was probed by independently tagging the protein with FLAG and HA tags (THAP9-F, THAP9-H) and observing if these differently tagged proteins colocalise and form a homodimer.

Immunoblot analysis established that THAP9-H and THAP9-F expressed the best 24 and 48 hours post transfection respectively (Fig. 1A). Interestingly, they were found to colocalise in the nuclei of HEK293T cells both 24h (Fig. 1B) and 48h post co-transfection (Fig. 1C). The colocalisation of THAP9-H and THAP9-F was statistically significant (P value < 0.0001, Fig. 1E) when compared to control (lipofectamine treated cells, Fig. 1D)

Since THAP9-H and THAP9-F showed optimal expression at different time points, pull-down assays were performed instead of co-transfecting both constructs followed by co-immunoprecipitation. When THAP9-F was used as a bait, it could pull down THAP9-H (Fig. 1F, lane 1). THAP9-H bound to beads was run as a positive control (Fig. 1F, lane 2).

Thus, colocalisation and pull-down experiments establish that Human THAP9 appears to undergo homo-oligomerisation in the nucleus.

### Prediction of amino terminal leucine rich coiled coil regions in DmTNP and hTHAP9

We have previously identified an evolutionarily conserved, leucine rich, alpha helical coiled coil region, downstream of the amino terminal THAP domain of most human THAP proteins including THAP9 (27). Since coiled coil regions mediate protein oligomerization (33), it was predicted that the identified coiled coil in human THAP9 may be involved in oligomerisation.

DmTNP, which is homologous to hTHAP9, is known to exist in different oligomeric forms at different steps of transposition; as a tetramer bound to DNA in the PEC/CDC (15) and as a dimer in the post-transposition STC (17). However, the recent cryoEM structure of the dimeric DmTNP STC could not resolve the flexible amino terminal DNA binding domain and adjacent dimerization domain (residues 1-150) (17) and thus the role of these domains in DmTNP oligomerisation is not clear. Hence we decided to probe if there was an additional oligomerisation region in the amino terminal region of DmTNP.

COILS analysis predicted that multiple short regions throughout the length of DmTNP had a higher probability (>0.5) of forming coiled coils (Fig. 2A, top panel). In hTHAP9, some short regions in the centre were predicted to have higher probability (>0.5) while the amino terminal region had modest probability (>=0.3) of forming coiled coils (Fig. 2A, top panel). JPRED predicted that these putative coiled coil regions had a very high tendency of forming alpha helices till residue 140 for DmTNP and till residue 176 for hTHAP9 (Fig. 2B). Interestingly, a strong alpha helical region was independently predicted till residue 165 in hTHAP9.

**Figure 2.**
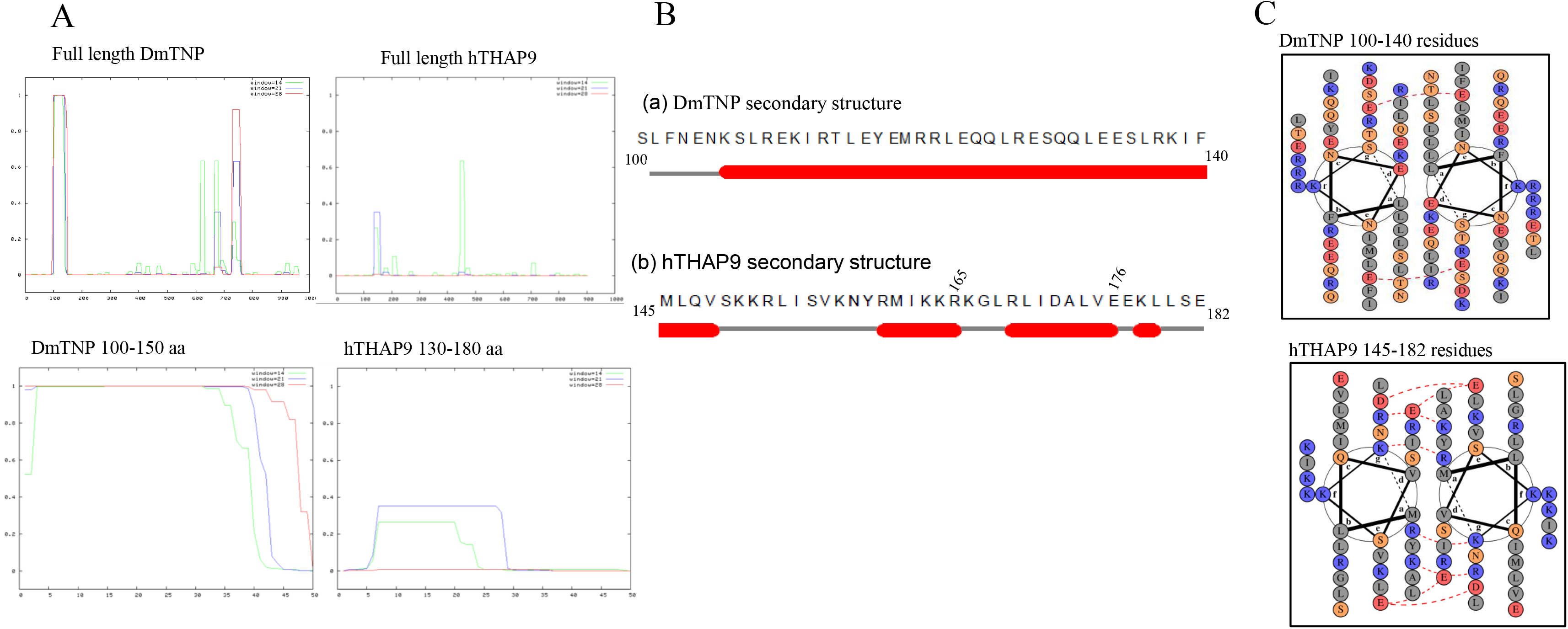
Prediction of amino terminal coiled coil region in DmTNP and hTHAP9. A. Top panel: COILS predicted regions with high probability of forming coiled coil structures in the DmTNP and hTHAP9, considering a window size of 14 (green curve), or 21 (blue curve) or 28 (red curve). Bottom panel: Zoomed view of amino terminal regions with high probability of forming coils (DmTNP:100-150 residues, hTHAP9: 130-180 residues) to define approximate boundaries. B. JPRED predicted secondary structures of the amino terminal coiled coil region. The red barrels represent alpha helices and the green arrow represents a beta sheet. C. DRAW COIL predicted electrostatic interactions in the predicted coiled coil regions represented as a helical wheel plot. Non-polar (grey), polar (yellow), acidic (red) and basic (blue) residues are marked. The red dotted lines represent electrostatic repulsive interactions and the blue dotted lines represent electrostatic attractive interactions

Coiled coils are characterised by “heptad repeats” (repeated pattern of seven amino acids, a-g), in which the *a* and *d* positions are non-polar/hydrophobic residues (like Leu) while the *e* and *f* positions are charged amino acids (Suppl. Fig. 2A). During protein oligomerisation, the side chains of hydrophobic residues on one monomer mediate ‘knob into holes’ packing with similar regions from another monomer. This leads to the formation of homo-oligomers (33), characterized by amphipathic alpha helices of individual monomers twisting around each other and enclosing a hydrophobic core. The regions immediately after DBD till the end of the predicted coiled coil region (hTHAP9: 90-165 residues; DmTNP: 78-140) were aligned to the heptad repeat pattern. Interestingly, some leucines in both proteins aligned to the *a* and *d* positions (Suppl. Fig. 2B, 2C) (27), consistent with previous reports (19, 27).

To investigate if the predicted alpha helical regions (from Fig. 2A; DmTNP: residues 100-140 and hTHAP9: residues 145-182) could form amphipathic helical structures via electrostatic interactions amongst themselves, DRAW COIL analysis was performed. Multiple electrostatic repulsive interactions were predicted in both of these regions as seen in Fig. 2C.

### Nuclear expression of truncated DmTNP and hTHAP9

Based on our bioinformatic analyses (COILS, JPRED, DRAW COIL; Fig. 2), we decided to experimentally investigate the role of the amino terminal DBD and leucine rich coiled coil regions in DmTNP (D=residues 1-140) and hTHAP9 (H1=residues 1-176 and H2=1-165). We constructed both HA- and FLAG-tagged carboxy-terminal truncated clones which included the amino terminal DBD (DmTNP- 1-77 residues, hTHAP9-1-94 residues), the predicted nuclear localization signal (NLS) (DmTNP- 60-69 residues, hTHAP9- 150-168 residues) and the predicted coiled coil regions [DmTNP-100-140 residues, hTHAP9- 145-165 (H2) or 145-176 (H1) residues] of each protein.

Each FLAG-tagged (F-H1, F-H2, F-D) and HA-tagged (H1-H, H2-H, D-H) truncated protein was individually found to express and localize to the nucleus of HEK293T cells (Fig. 3A and 3B). HEK293T cells treated with lipofectamine alone (bottom panels) confirm that there was no background signal. Western analysis demonstrated that all these constructs [approximate MW: 20 kDa (H1), 19 kDa (H2), 17 kDa (D)] expressed optimally 48h post-transfection (Fig. 3C and 3D).

**Figure 3.**
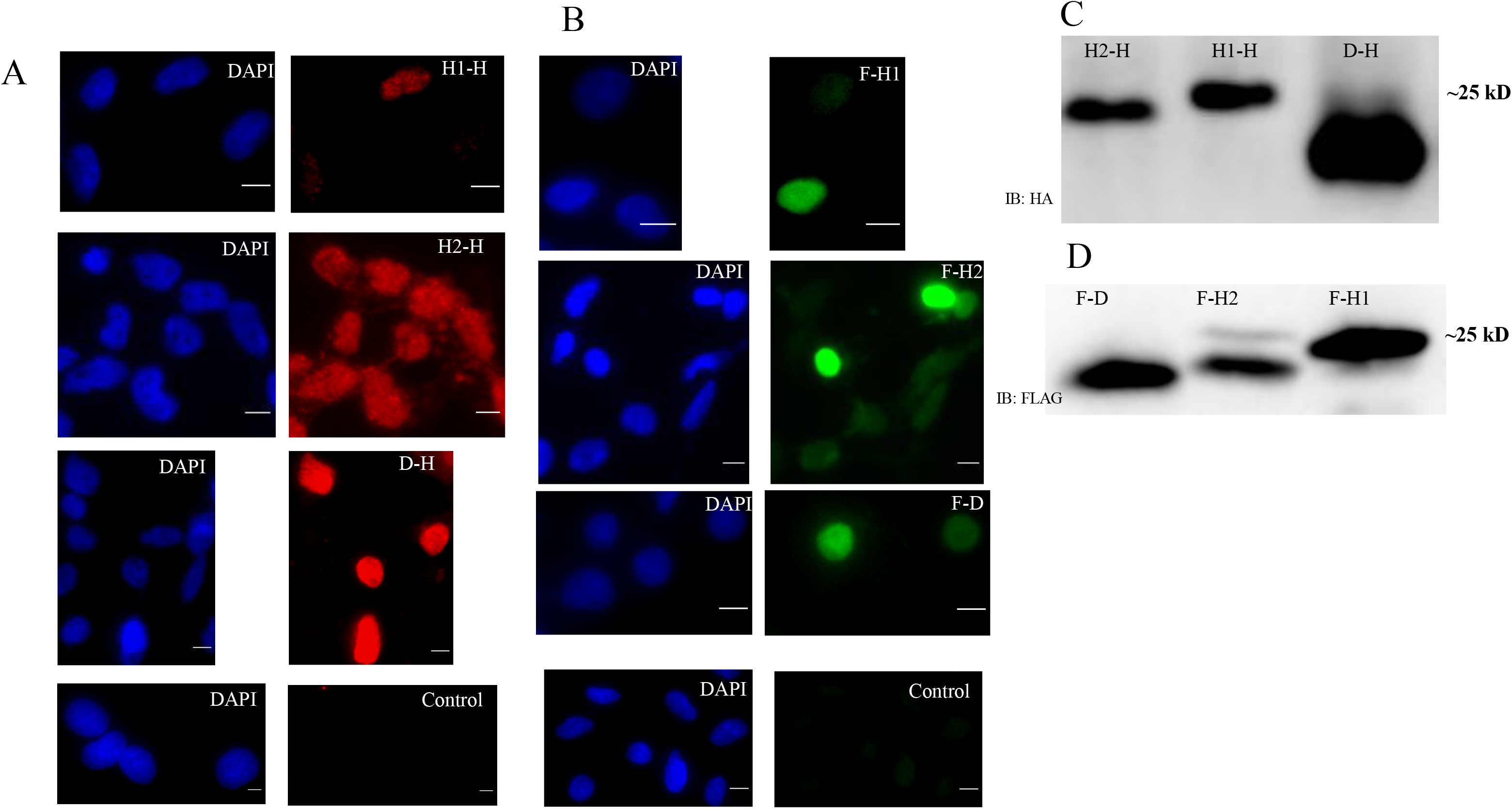
Nuclear expression of truncated DmTNP and hTHAP9. HEK293T cells were individually transfected with (A) HA-tagged constructs (H1-H, H2-H, D-H) (B) FLAG-tagged constructs (F-H1, F-H2, F-D) and immunostained for FLAG (green) and HA (red). Control (lipofectamine only), nuclei were counterstained with DAPI (blue). The images shown here were captured by Leica confocal microscope (10× eyepiece and 40× objective). Scale bars= 10 microns. Immunoblotting whole cell lysates of 293T cells harvested 48 hrs after being transfected with (C) H1-H, H2-H, D-H (anti-HA) (D) F-H1, F-H2, F-D (anti-FLAG).

### Nuclear co-localisation of truncated DmTNP and hTHAP9 suggests homo-oligomerisation

Truncated differentially tagged DmTNP (F-D, D-H) or hTHAP9 (F-H2, H2-H or F-H1, H1-H) were found to co-localise in the nucleus. Briefly, immunostaining of HEK293T cells co-transfected with either F-H1 and H1-H (Fig. 4A), F-H2 and H2-H (Fig. 4B) or F-D and D-H (Fig. 4C) demonstrated that truncated DmTNP or hTHAP9 appear to homo-oligomerise in the nucleus. The colocalisation was statistically significant with P value < 0.0001 (Fig. 4E).

**Figure 4.**
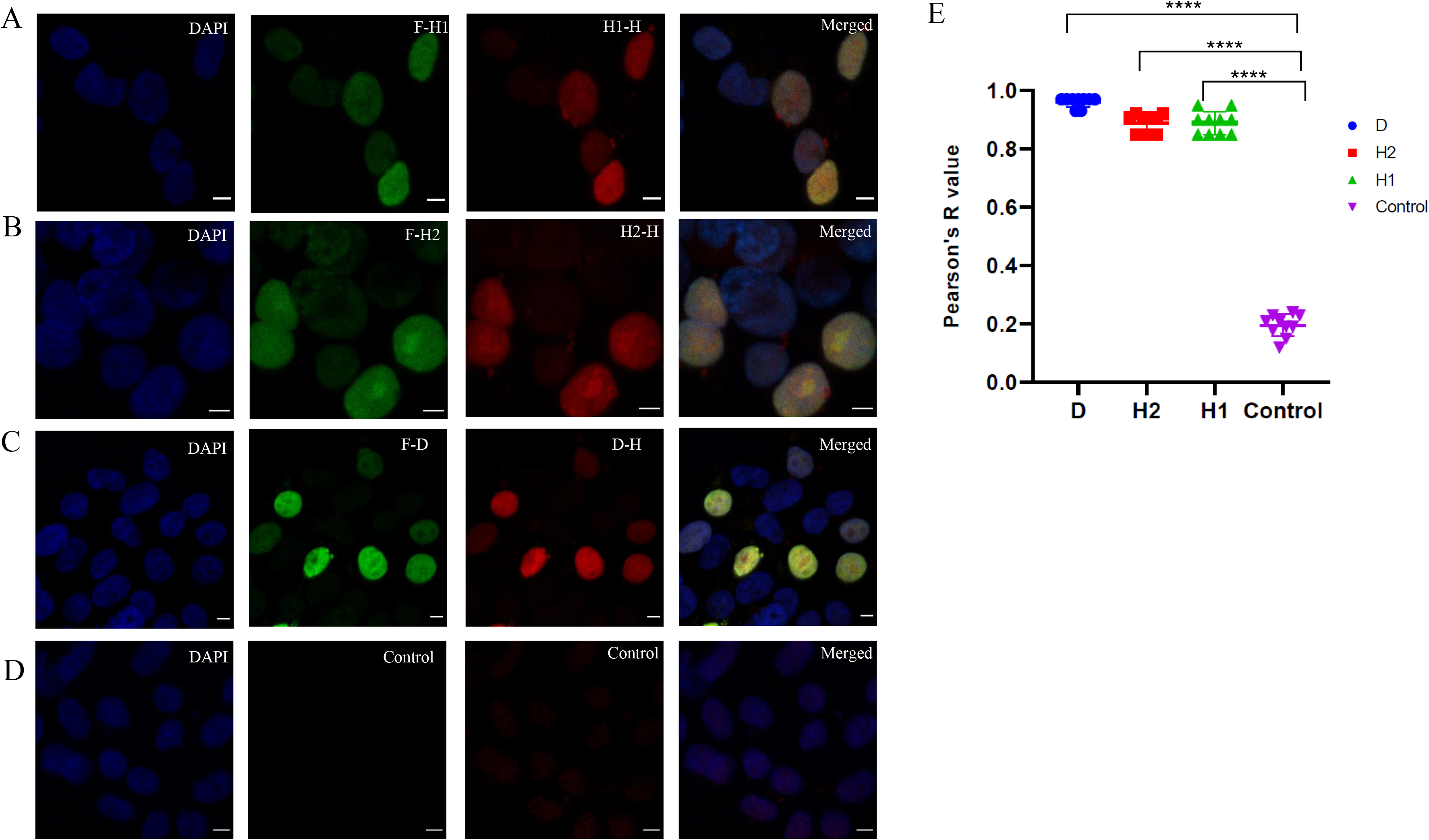
Nuclear co-localisation of truncated differentially tagged DmTNP (F-D, D-H) and hTHAP9 (F-H2, H2-H or F-H1, H1-H) HEK293T cells co-transfected with (A) F-H1 and H1-H (B) F-H2 and H2-H (C) F-D and D-H (D) Control (lipofectamine only) were immunostained for FLAG (green) and HA(red). Nuclei were counterstained with DAPI (blue). The images shown here were captured by Leica confocal microscope (10× eyepiece and 40× objective). Scale bars= 10 microns. (E) Pearson’s R-value (no threshold) for co-localisation of constructs corresponding to panels 4A, 4B or 4C compared to 4D. The number of cells counted for each sample n=10.

The observed nuclear co-localisation may be due to the spatial overlap or proximity of these truncated proteins in the nucleus. To further probe this, we performed proximity ligation assays (PLA), which can detect possible interactions between two proteins, less than 40 nm apart, as foci (red). Briefly, HEK293T cells co-transfected individually with (Fig. 5A: F-H1 and H1-H; F-H2 and H2-H or F-D and D-H) were fixed and used for PLA using two different antibody pairs. Counting PLA foci (red) established that F-H1 and H1-H; F-H2 and H2-H as well as F-D and D-H occur in close proximity (P <0.0001) when compared to control (lipofectamine treated HEK293T cells).

**Figure 5:**
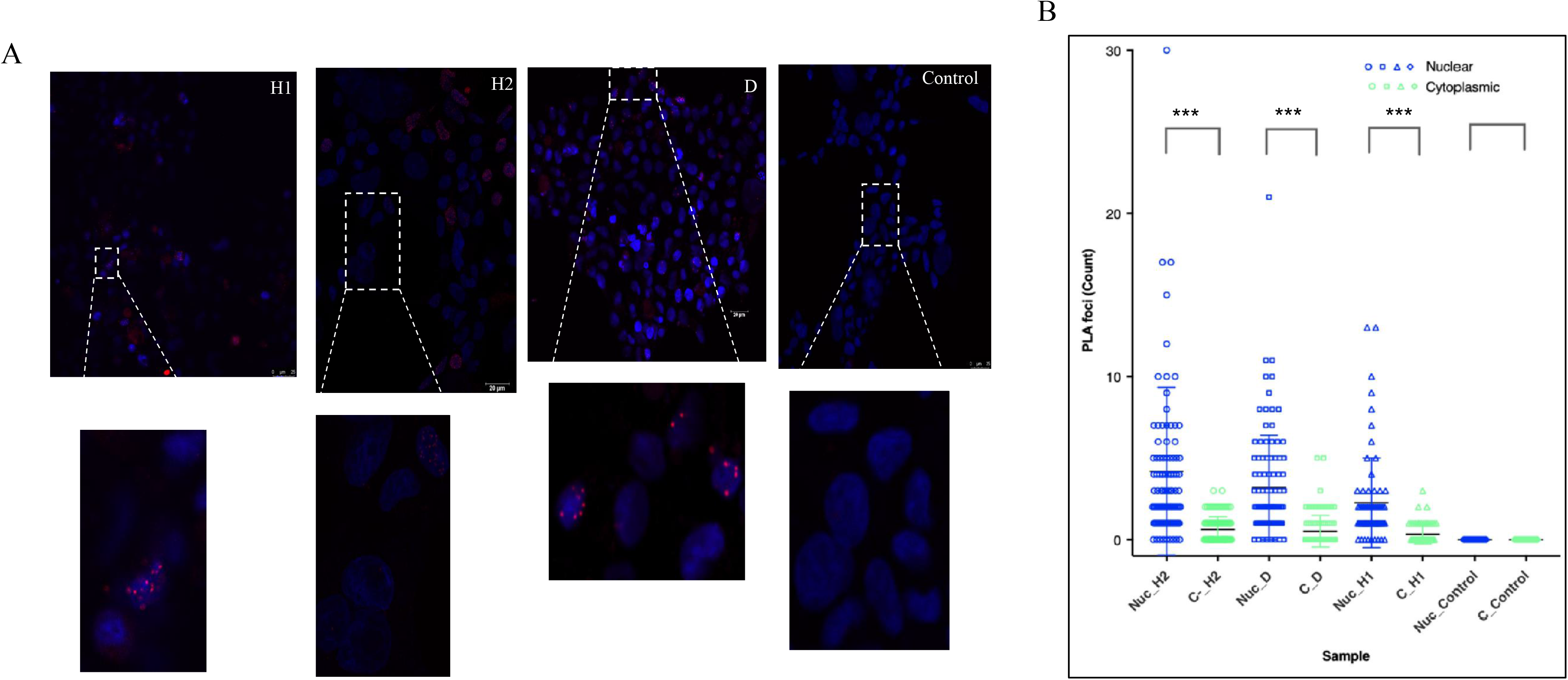
The proximity of truncated DmTNP and hTHAP9 interactions were stronger in nuclei. (A) HEK293T cells co-transfected individually with (F-H1 and H1-H, F-H2 and H2-H and F-D and D-H, lipofectamine alone) were fixed and used for PLA. The nuclei were counterstained with DAPI (blue) Insets in lower panel focus on PLA foci in nuclei (B) Comparison of the PLA signals in the nucleus (blue) and cytosol (green) of each truncated construct and the control. The images shown here were captured by Leica confocal microscope (10× eyepiece and 40× objective), scale bars= 10 microns. D, H2,control (n=100); H1(n = 70)

Although, some PLA foci were detected in the extranuclear region, (Fig. 5 A) H1, H2 and D homo-oligomerisation was more prevalent in the nuclei (P value <0.0001) than the cytoplasm. The preferred nuclear interaction could be due to the presence of the DBD and predicted NLS in the truncated constructs.

### Homo-oligomerisation of truncated DmTNP and hTHAP9 confirmed by co-immunoprecipitation

To further confirm if the observed co-localisation (Fig. 4 and 5) was due to the interaction of the truncated proteins, we performed reciprocal co-immunoprecipitation assays on whole cell lysates obtained from HEK293T cells co-transfected with F-H1 and H1-H, F-H2 and H2-H or F-D and D-H. Immunoprecipitation of F-H2 and F-D using HA antibody and the reciprocal immunoprecipitation of H2-H and D-H using FLAG antibody, confirmed the interaction between F-H2 and H2-H (Fig. 6A) and F-D and D-H (Fig. 6B).

**Figure 6:**
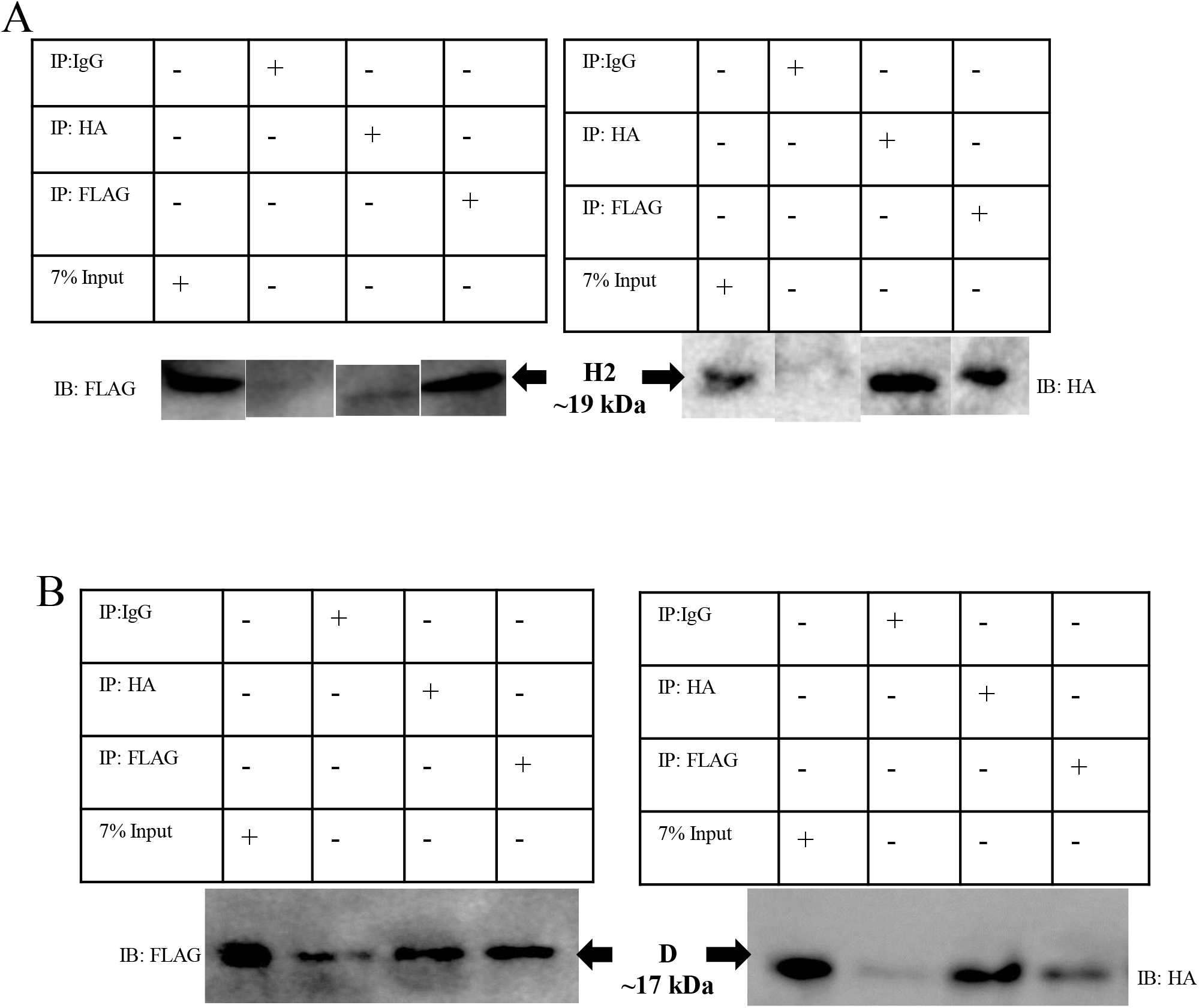
Co-immunoprecipitation of truncated DmTNP and hTHAP9 suggest their homo-oligomerisation. Reciprocal co-immunoprecipitation followed by immunoblotting of (A) H2-H and F-H2 (B) D-H and F-D. Western blot (left panel: anti-FLAG, right panel: anti-HA) IgG (negative control), input fraction (positive control)

Interestingly, the reciprocal co-immunoprecipitation of F-H1 and H1-H was very weak (Suppl. Fig. 3). H1 (residues 1-176) has 11 additional amino acids (residues 166-176) when compared to H2 (residues 1-165). Thus, this suggests that the 1st 165 residues of hTHAP9 (H2) may constitute a minimal amino-terminal oligomerisation domain. It is also intriguing to note that the addition of residues 166-176 significantly abrogate the oligomerisation, without affecting protein expression.

### Role of Leucines and oppositely charged amino acid pairs in oligomerisation

The predicted helical coiled coil regions of DmTNP and hTHAP9, which appear to be important for homo-oligomersation, are leucine rich (Fig. 2B, Suppl. Fig. 2).

DNA transposases such as IS911 have been reported to form multimers mediated by leucine rich coiled coil structures (7). The DmTNP inhibitor KP repressor has also been shown to dimerise via leucine rich coiled coil (19). Thus, we decided to investigate if the leucine rich regions (~40 residues long, Suppl. Fig.2) downstream of the amino terminal THAP domain in hTHAP9 (145-182 residues) (27) and DmTNP (100-140 residues) were involved in oligomerisation.

When the Leucine-rich regions of DmTNP and hTHAP9 were aligned to the heptad repeat pattern characteristic of coiled coils, several leucines in both proteins aligned to the *a* and *d* positions (Suppl. Fig. 2B, 2C). These residues (DmTNP: Leu at positions 101, 108, 115, 122, 132 and 136; hTHAP9: Leu at positions 90, 128, 132, 139, 146 and 153) were individually mutated to Ala (polarity and size similar to leucine) in the corresponding truncated proteins (D-H, H2-H respectively). The LtoA point mutants (on H2-H or D-H templates) were individually probed for disruption of interaction by reciprocal co-immunoprecipitation with F-H2 or F-D respectively. L90A_H2H and L101A_DH were not included in the co-IP experiments as they did not express well. Surprisingly, none of the LtoA point mutants of either hTHAP9 (Fig. 7A) or DmTNP (Fig. 7B) showed complete disruption of interaction. To check for the possible combined effect of all the Leucines, we created combined LtoA mutants for both hTHAP9 [H2(all)-H (L90A, L128A, L132A, L139A, L146A, L153A)] and DmTNP [D(all)-H (L101A, L108A, L115A, L122A, L132A, L136A)]. However, even the combined leucine mutants did not show complete disruption of interaction either for hTHAP9 (Fig. 7C) or for DmTNP (Fig. 7D).

**Figure 7:**
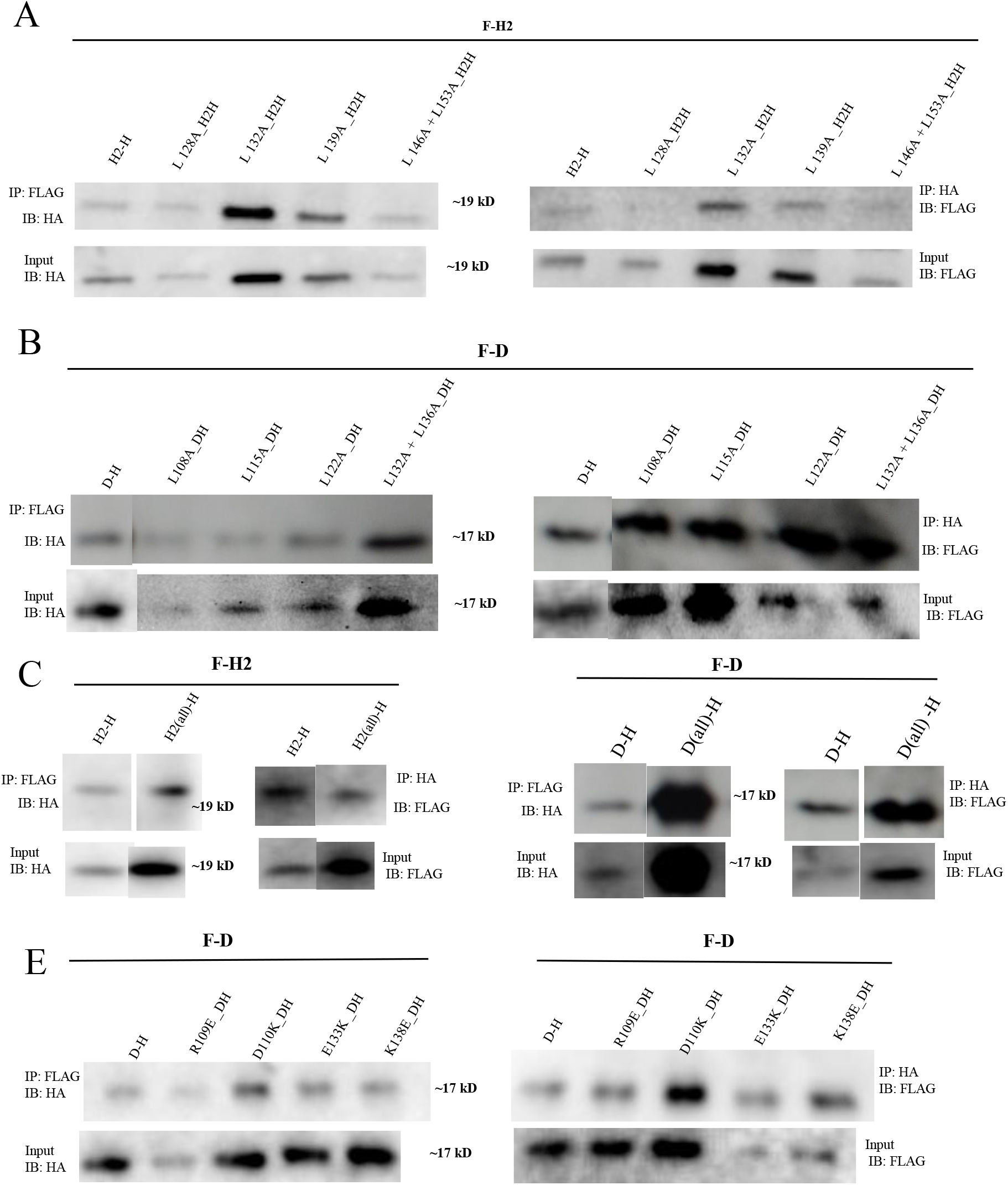
Leucine and charged pair mutations in truncated DmTNP and hTHAP9 does not disrupt homo-oligomerisation. Reciprocal co-immunoprecipitation (left panel; IP: anti-FLAG, right panel; IP: anti-HA) followed by immunoblotting (left panel: anti-HA, right panel: anti-FLAG) of (A) F-H2 with H2-H (wildtype) or each leucine point mutant in H2H (L128A_H2H, L132A_H2H, L139A_H2H) or a double leucine mutant (L146A+L153A_H2H) (B) F-D with D-H (wildtype) or each leucine point mutant in D-H (L108A_DH, L115A_DH, L122A_DH) or a double leucine mutant (L132A+L136A_DH) (C) Combined leucine mutant in hTHAP9; H2(all)-H (L90A, L128A, L132A, L139A, L146A, L153A) with F-H2 (D) Combined leucine mutant in DmTNP; D(all)-H (L101A, L108A, L115A, L122A, L132A, L136A) with F-D (E) Each charged pair point mutant in DmTNP (R109E_DH, E110K_DH, E133K_DH, K138E_DH) with F-D. The expression of each construct was shown by immunoblotting input (7%) with anti-FLAG (right panel) and anti-HA (left panel) (panel below respective immunoprecipitation fractions)

It has previously been reported that double mutants of KP repressor protein (first 199 residues identical to DmTNP), in which L108 and L115 were mutated to either alanine, valine, aspartic acid and proline, retained the ability to dimerise in vitro (19). In a separate study it was found that mutating L101 and L122 in KP repressor protein, to either isoleucine and arginine or valine and histidine respectively, did not completely disrupt its ability to repress P element transposition in fruit flies (20).

NMR studies on THAP11 demonstrated that its carboxy terminal region (254-306 residues) formed a homodimer, which was partly stabilized by a salt bridge interaction between an oppositely charged pair (K299 and D300) that occurred at positions *e* and *g* of its heptad repeat pattern (34). Thus, we decided to investigate if the acidic-basic residue pairs (Suppl Fig. 2C, green residues) in DmTNP (R109 and E110, E133 and K138) played any role in its oligomerisation. Acidic residues E110 and E133 were mutated to basic residues i.e. Lys (E110K, E133K) while basic residues R109 and K138 were mutated to acidic residues i.e.Glu (R109E, K138E) in truncated DmTNP (D-H). However, it was observed that none of these point mutations (R109E, D110K, E133K and K138E) disrupted the interaction with F-D in reverse co-IP experiments (Fig. 7E).

Thus, leucine and charged pair mutations in truncated DmTNP and hTHAP9 does not disrupt homo-oligomerisation

### Truncated DmTNP and hTHAP9 can oligomerise without the coiled coil region

The leucine residues in the coiled coil region did not appear to be important for hTHAP9 and DmTNP oligomerisation. Thus, we decided to delete the predicted oligomerisation region (DmTNP: 100-140 residues, hTHAP9: 145-165 residues) in both proteins to ascertain its importance. We constructed HA-tagged truncated clones which included the first 144 residues of hTHAP9 (H_Del-H) and the first 99 residues of DmTNP (D_Del-H;1-99 residues): both these constructs included the amino terminal DBD (DmTNP- 1-77 residues, hTHAP9- 1-89 residues) of each protein.

In a previous study, in vitro chemical cross-linking experiments using purified proteins reported that shorter DmTNP constructs containing the first 88 or 98 residues were unable to dimerise (18, 19). However, surprisingly, reciprocal co-immunoprecipitation demonstrated that neither H_Del-H (Fig. 8A) nor D_Del-H (Fig. 8B) lost the ability to homo-oligomerise. Thus, it appeared that both DmTNP and hTHAP9 were able to oligomerise in cells even without the predicted Leu-rich coiled region. More importantly, these surprising results suggested that the DBD and the region between the DBD and predicted coiled coil region may help mediate oligomerisation.

**Figure 8.**
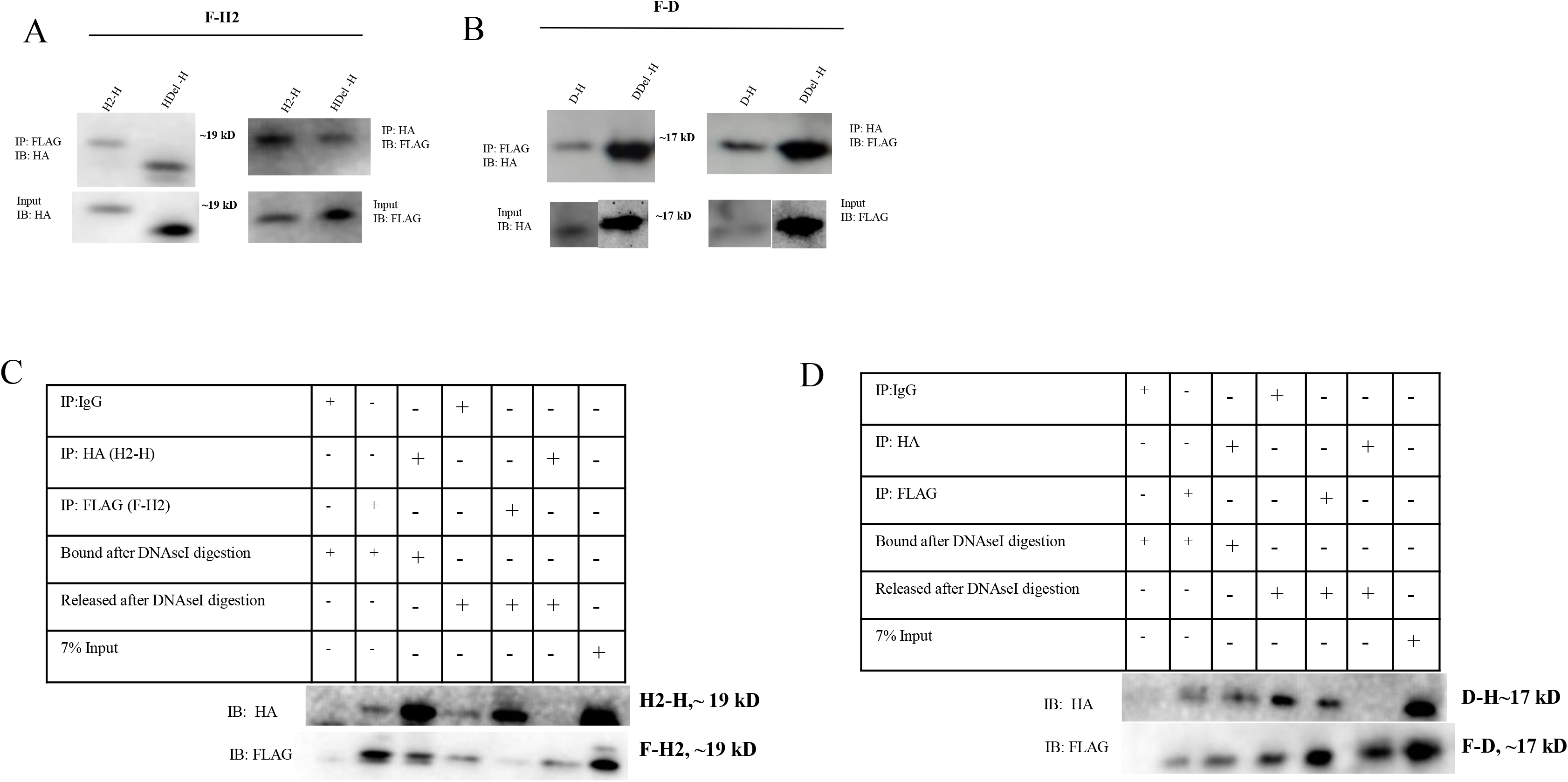
Interaction between the truncated DmTNP and hTHAP9 constructs is partially mediated by DNA. Reciprocal co-immunoprecipitation followed by immunoblotting (left panel: anti-HA, right panel: anti-FLAG) of (A) hTHAP9 deletion mutant (Hdel-H;1-144 residues) with F-H2 (B) DmTNP deletion mutant (DDel-H;1-99 residues) with F-D. The expression of each construct was shown by immunoblotting input (7%) with anti-FLAG (right panel) and anti-HA (left panel) (panel below respective immunoprecipitation fractions) Reciprocal co-immunoprecipitation followed by cleavage with DNaseI on beads bound to the (C) immunoprecipitated H2 homodimer (F-H2 and H2-H) as seen in both the beads-bound fraction and released-from-beads fraction. (D) immunoprecipitated D homodimer (F-D and D-H) as seen in both the beads bound fraction and the released from beads fraction.

### DNA mediates interaction between truncated DmTNP and hTHAP9

DNA transposases such as MuA and *Hermes* have been reported to form DNA-mediated homo-oligomers (8,9). Our surprising observations that both H_Del (Fig. 8A) and D_Del (Fig. 8B) were able to interact with F-H2 and D-H respectively, led us to hypothesize that these interactions could be mediated by DNA.

To investigate this further, on-beads DNaseI cleavage assays (Suppl. Fig. 4) were performed after reciprocal co-immunoprecipitation of F-H2 and H2-H (Fig. 8C) or F-D and D-H (Fig. 8D). If DNA mediates the interactions between each set of co-immunoprecipitated monomers bound to beads, the proteins should not remain bound to beads after DNaseI cleavage, as seen in Fig. 8C and 8D. On the other hand, if homo-oligomerisation is not DNA-mediated, DNase cleavage will not release the bound protein from beads. It is to be noted that DNase cleavage did not release all the bound protein (Lanes 2 and 3 in Fig. 8C and 8D); this may be due to incomplete DNase cleavage because of inaccessibility of the bound proteins to the enzyme or because the observed homo-oligomerisation of hTHAP9 and DmTNP truncated proteins is partially DNA mediated.

### Human THAP9 may interact with Hcf-1 and other human THAP proteins

Having established that hTHAP9 is able to homo-oligomerise, we decided to explore if the protein could also undergo hetero-oliogomerisation. Identifying hTHAP9’s protein interaction partners would also help understand its yet unknown cellular function. Interestingly, the STRING database (24) predicted that some human THAP family members namely THAP1, THAP10 and THAP11 may interact with hTHAP9. Moreover, hTHAP9 harbors a predicted Hcf-1 binding consensus motif [HBM, (D/E)HXY)] (25) between 123-126 residues. Thus, we decided to investigate if hTHAP9 could interact with Hcf1, THAP1, THAP10 and THAP11.

THAP9-F was found to colocalise with H-Hcf-1 48h post transfection (Fig. 9A). Also, THAP9-H was found to individually co-localize with THAP1-F (Fig. 9B), F-THAP10 (Fig. 9C) and F- THAP11 (Fig. 9D) 24h post transfection. The co-localization of THAP9 with other protein interaction partners was found to be statistically significant (P value < 0.0001) when Pearsons’s R-value was compared to that of the control, as shown in Fig. 9E.

**Figure 9.**
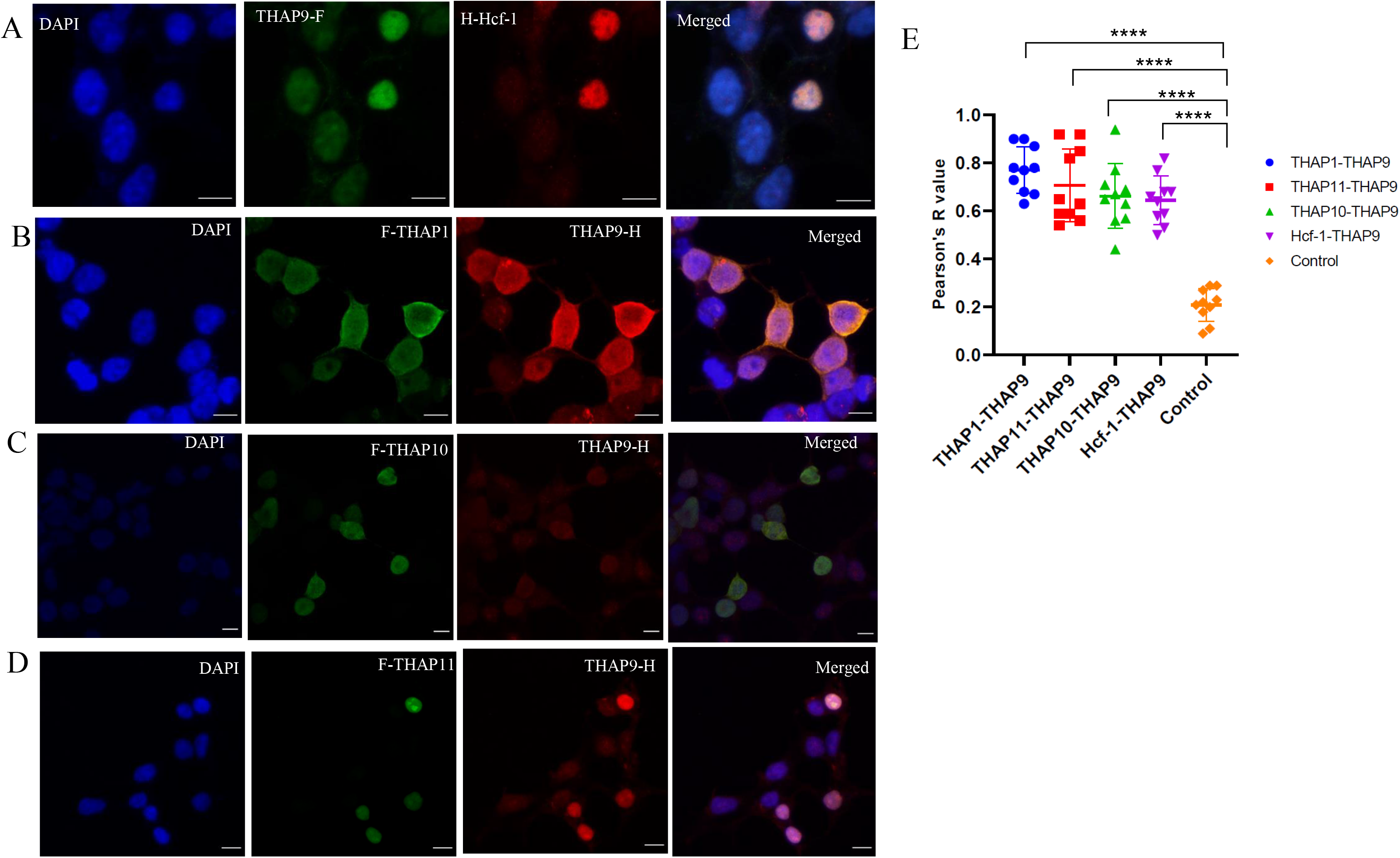
hTHAP9 colocalises with hTHAP1, hTHAP10, hTHAP11 and human Hcf-1 in the nucleus of HEK293T cells. HEK293T co-transfected with (A) H-Hcf-1 and THAP9-F, (B) THAP9-H and THAP1-F (C) THAP9-H and F-THAP10 (D) THAP9-H and F-THAP11 were immunostained for FLAG (green) and HA (red). Nuclei were counterstained with DAPI (blue). (E) Comparison of Pearson’s R-value (no threshold) for co-localization of of constructs corresponding to panels 9A, 9B. 9C and 9D with the control (lipofectamine treated HEK293T cells). The number of cells counted for each sample n=10. The images shown here were captured by Leica confocal microscope (10× eyepiece and 40× objective). Scale bars= 10 microns.

To define the minimal region required by hTHAP9 to interact with some of its partners, we checked if THAP1 or THAP10 could interact with truncated hTHAP9 (H2). Interestingly, H2-H did not co-immunoprecipitate with either THAP1-F or F-THAP10 (data not shown). This indicates that in addition to the amino terminal interaction regions described in this study, hTHAP9 may have additional carboxy terminal interaction domains which interact with THAP1 and THAP10.

Overall, we demonstrate that hTHAP9, like its homolog DmTNP, can undergo homo-oligomerisation, which is partially mediated by DNA. Deletion mutant, leu, H1Moreover, hTHAP9 can also hetero-oligomerise with several members of the human THAP family as well as Hcf1.

## Discussion

Protein oligomerisation, wherein two or more polypeptide chains/subunits associate with each other, occurs across many protein families like bZIP transcription factor family (35), DNA transposases (8), Hsp70 (36) and GADs (GTPases activated by Dimerization) (37). This process can offer many functional advantages like evolution of functional control or higher-order complexity (38). The strength and duration of protein oligomerisation is often regulated by various parameters like protein concentration, temperature, pH, binding of ligand, nucleic acid or nucleotides, phosphorylation state, etc.(38)

The homo oligomerisation of DmTNP observed in our results is consistent with earlier reports of DmTNP forming functional oligomers [tetrameric in early steps of transposition (15) and dimeric in the post transposition transpososome complex (17)]. The internally deleted 24kD KP repressor (18) may form a heteromultimeric complex with DmTNP (20). Moreover, the 66kD Type I repressor protein, encoded by an alternatively spliced P element isoform, represses somatic P element transposition (5). It is exciting to speculate that the 66kD protein, the first 561 amino acids (including the DBD and predicted coiled coil region) of which are identical to DmTNP, may also associate with active DmTNP to form inactive multimers.

There have been no previous reports of oligomerisation of the DmTNP homolog, hTHAP9. This study, reports for the first time, that hTHAP9 too, can undergo homo-oligomerisation. We also predict that a leucine rich, alpha helical coiled coil region immediately downstream of the DBD may be partially important for protein-protein interactions. Since the cellular role of hTHAP9 is still unknown, it is exciting to speculate that the observed oligomerisation may have important functional consequences ranging from aiding DNA binding, excision and/or integration, recruitment of other proteins which then signal a downstream cascade of events etc.

It is to be noted that mutating the leucines (either individually or together) or deleting the Leucine-rich predicted coiled coil region, in both DmTNP and hTHAP9 did not disrupt homo-oligomerisation. This observation is in contrast to reports of leucine zipper mediated oligomerisation of proteins [e.g., AKAP-Lbc (39)]. On the other hand, the significant disruption of hTHAP9 oligomerisation by the presence of residues 166-176 (Suppl. Fig 3) in the truncated construct, suggest that there may be certain surfaces that inhibit association. The observation, that both DmTNP and hTHAP9, can oligomerise without the predicted coiled coil region also suggests that both these proteins have multiple putative oligomerisation regions. This hypothesis is supported by the recent cryoEM structure of DmTNP which shows that although the flexible amino terminal 150 residues was not clearly resolved, the rest of the protein (151-751 residues) dimerised via several DNA-based interactions throughout the interlocked protein. Notably, a single DmTNP subunit catalyzes strand transfer (via its RNaseH and GTP-binding domains) of one transposon DNA end but binds the other transposon end via its HTH and C-terminal domains, thus supporting catalysis in *trans* (17).

We also report a novel finding that hTHAP9 and DmTNP oligomerisation may be partially mediated by DNA. DNA dependent oligomerisation has been reported for Tn5 and MuA transposases (8). *In vitro* experiments using purified protein have previously shown that mutations in the C2CH motif of the THAP domain, abolishes binding to specific P element DNA sites but not dimerization of the KP repressor protein (18, 19). However, both Fig.8C and 8D demonstrate that in the cellular context, homo-oligomerisation of truncated hTHAP9 and DmTNP is partially mediated by genomic DNA. This can be explained by the possibility of hTHAP9 and DmTNP directly interacting with human genomic DNA, non-specifically. Alternately, these proteins may indirectly interact with DNA, by being associated with other partners that are directly bound to DNA.

It would be interesting to study the role of DBD (DmTNP: residues 1-77, hTHAP9: residues 1-94) in oligomerisation and the adjacent leucine rich coiled coil domain in binding DNA. Previous *in vitro* studies of DmTNP, have shown that shorter constructs containing the first 88 or 98 residues were unable to dimerise (19) while the DBD alone could bind P element DNA with high affinity (40). Further studies would pave the way for understanding whether for hTHAP9, DNA binding and oligomerisation are mutually dependent activities or if there is a particular order of reactions i.e. oligomerisation followed by DNA binding, as shown in DmTNP (15) or vice versa.

We also report that hTHAP9 and both truncated hTHAP9 and DmTNP (which include the NLS) express preferentially in the nucleus of HEK293T cells. However, the functional implications of nuclear localization is yet to be investigated. It is interesting to note that the nuclear import of *Mos1* Mariner transposase is essential for transposition and is dependent on Mos1’s NLS and dimerisation domain (41). It is tempting to speculate that hTHAP9 and DmTNP may function similar to Mariner transposase.

The exact stoichiometry of hTHAP9 oligomeric state is however not clear. Does it form dimers, tetramers or higher order oligomers like octamers? Like DmTNP, does hTHAP9 exist in different oligomeric forms depending on the step of transposition? It will be interesting to explore the possible relationships between the oligomeric state/s of the protein and its corresponding functional status.

DmTNP binds GTP via an identified GTP binding region (G domain) as a prerequisite to forming the tetrameric synaptic paired end complex (PEC) bound to both ends of transposon DNA (16). There are no reports of GTP hydrolysis by DmTNP. hTHAP9, too, has a predicted G domain, although it is not known if it can bind or hydrolyse GTP. It is interesting to speculate that DmTNP (and possibly hTHAP9) are GADs (GTPases activated by Dimerization). GADs are known to adopt an active functional state after homo-dimerisation via GTP-bound G domains of individual monomers and their biological activity is terminated by GTP hydrolysis, which dissociates the functional dimer (37). The truncated proteins used in this study lack both the GTP binding and catalytic domains found in full-length DmTNP or hTHAP9. Thus, although the amino-terminal region of the transposase has been shown to mediate protein oligomerization, it is possible that the GTP binding and/or catalytic domains might mediate higher order oligomerization, important for assembly of an active transposase multimer during transposition.

Lastly, we have identified THAP1, THAP10, THAP11 and Hcf-1 as potential interaction partners of hTHAP9 (Fig. 9 B,C,D). The cellular roles of THAP1 include functioning as a transcription factor (42), regulation of endothelial cell proliferation (43), functioning as nuclear pro-apoptotic factor as well as transcriptional regulation at the inception of myelination in the oligodendrocytes (44). THAP10 is a less studied protein which was recently reported to be a downstream target of miR-383 that plays a role in inhibiting proliferation but promoting differentiation of Acute Myeloid Leukemia (AML) cells that harbour translocation between chromosome 8 and chromosome 21 (t(8;21)) (45). THAP11 is a transcription factor which governs the development of heart (46), retina (47) and brain (48) in addition to its demonstrated roles in pluripotency (49–51) and hematopoiesis (52).

Hcf-1 (Host Cell Factor) is a key regulatory protein involved in diverse cellular pathways like cell cycle progression, embryonic stem cell pluripotency and stress response (26) Hcf-1 is reported to interact with THAP1 and THAP3 using the amino terminal Kelch domain (3-455 residues). THAP1 is essential for recruitment of Hcf-1 during endothelial cell proliferation (43) while THAP3 was reported to enable Hcf-1 recruitment to promoters for transcriptional activation (25). Also, THAP11 is found to mediate Hcf-1 recruitment to the promoters of E2F target genes (53).

Since the exact cellular function of hTHAP9 is not known, it is exciting to speculate that given this association, hTHAP9 may be involved in one or many of the cellular pathways and processes in which THAP1, THAP10, THAP11 and Hcf-1 participate.

Of the twelve human THAP family proteins (THAP0-THAP11), THAP0 (54), THAP1 (55), THAP7 (56) and THAP11 (34) are reported to form homo-oligomers. Some THAP proteins also hetero-oligomerise in order to perform their cellular functions. For example, THAP0 forms a hetro-oligomer with MST1 (54), THAP3 associates with *O*-GlcNAc transferase (OGT) (25), THAP7 interacts with TAF-Iβ (56), histone H4 and HiNF-P (57) and THAP11 interacts with PolyC-binding protein 1 (PCBP1) (58). Thus, it is interesting to note that THAP9, like other members of the human THAP family, has been demonstrated to undergo both homo- and hetero-oligomerisation. Further studies are required to identify additional hTHAP9 interaction partners and establish their roles in the cellular and physiological functioning of hTHAP9.

Overall, we demonstrate that hTHAP9, like its homolog DmTNP, can undergo homo-oligomerisation, which is partially mediated by DNA and the amino-terminal end of the proteins. Moreover, hTHAP9 can also hetero-oligomerise with several members of the human THAP family as well as Hcf1.

## Supplementary figures

**Suppl. Figure 1.**
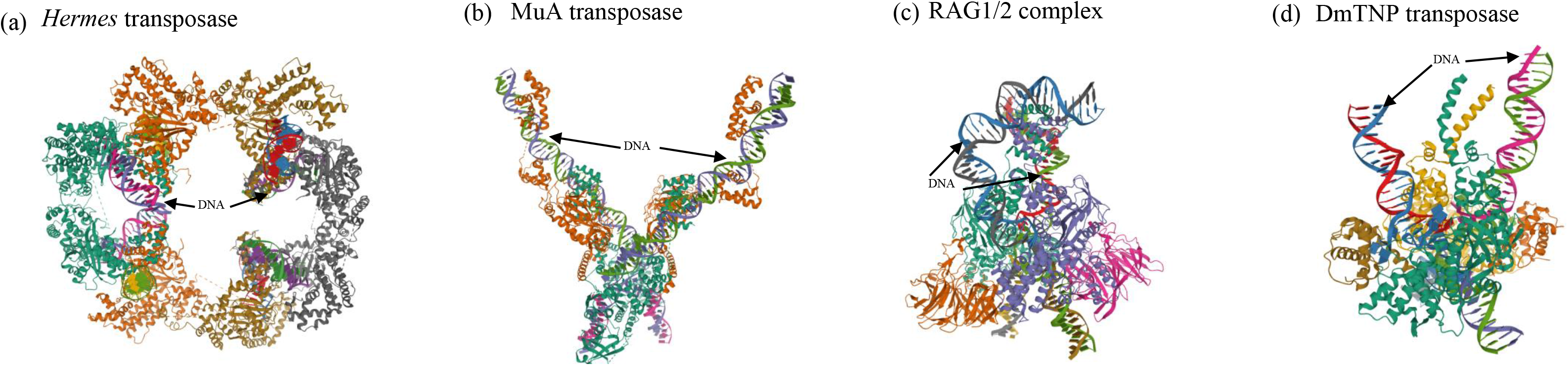
Oligomeric forms of different active DNA transposases and TE derived proteins. (a) *Musca domestica Hermes* forms an octamer (pdb id: 4D1Q) (b) Bacteriophage MuA forms a dimer in the presence of DNA (pdb id: 4FCY)(c) mouse RAG1/2 forms a heterotetramer (pdb id: 6OET) and (d) DmTNP forms a dimer in the presence of DNA and GTP (pdb id: 6PE2).

**Suppl. Figure 2.**
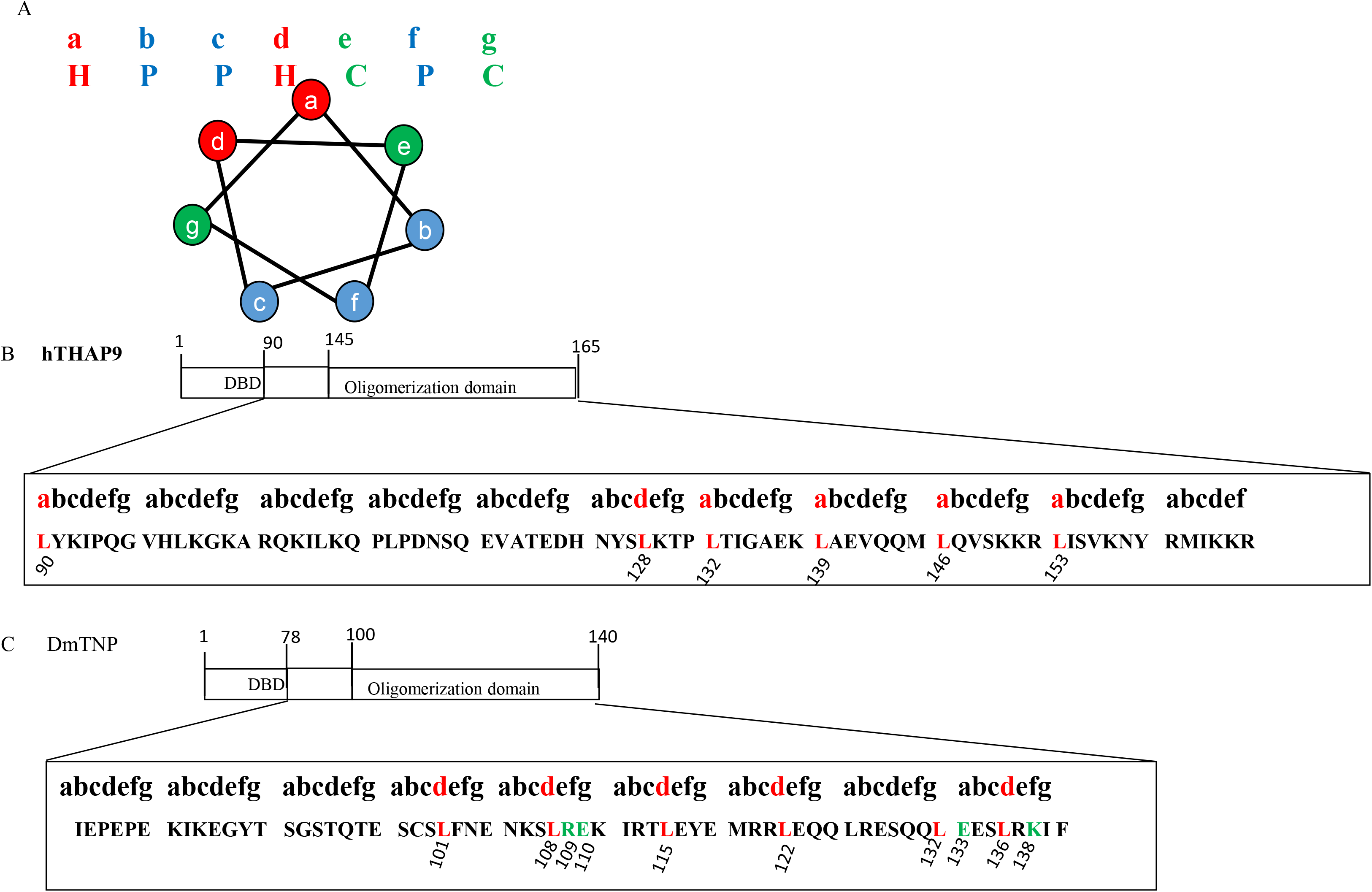
Alignment of the predicted coiled coil regions of hTHAP9 and DmTNP to the heptad pattern. (A) A typical heptad pattern of a coiled coil containing amphipathic arrangement of amino acids. The heptad ranges from *a* to *e* where a consensus arrangement includes hydrophobic amino acids (H; red) at positions *a* and *d*, polar amino acids (P; blue) at positions *b, c* and *f* and charged amino acids (C; green) at positions *e* and *g*. Leucines in hTHAP9 (B) and DmTNP(C) immediately after DBD and in predicted coiled coil region (B: 90-165 residues; C: 78-140) align with *a* and *d* (marked in red) of the heptad pattern. Two pairs of acidic and basic residues in DmTNP (R109 and E110, E133 and K138) are marked in green.

**Suppl. Figure 3.**
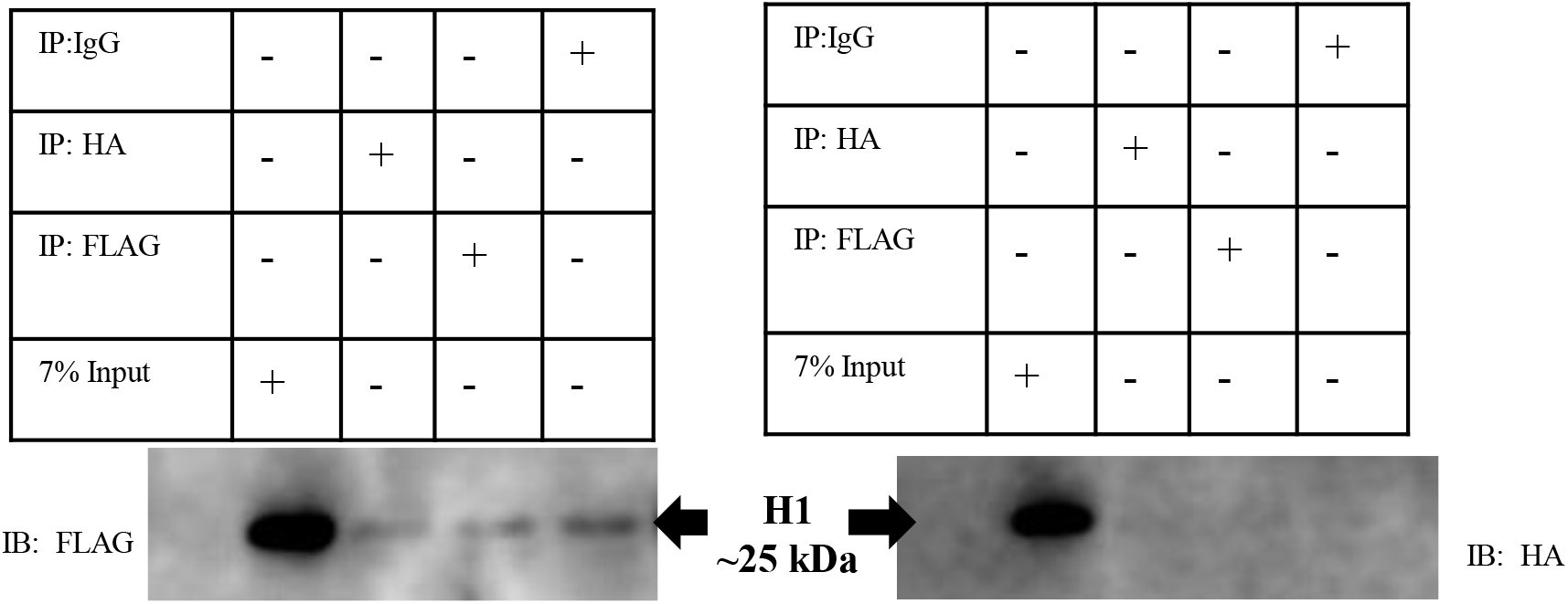
Co-immunoprecipitation of FLAG-H1 and H1-HA. Reciprocal co-immunoprecipitation of F-H1 and H1-H followed by Western with (A) anti-FLAG (B) anti-HA antibodies.

**Suppl. Figure 4.**
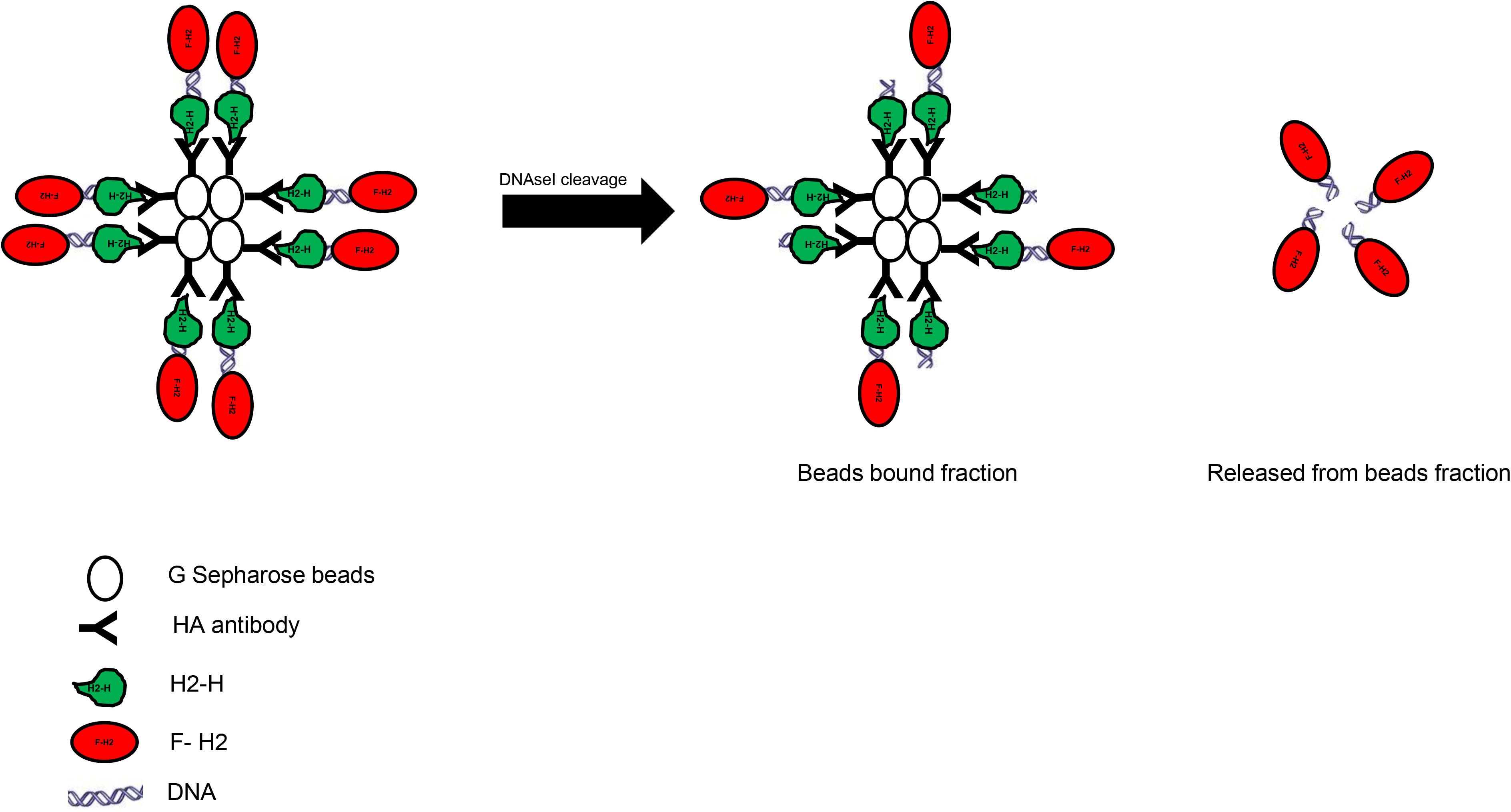
Schematic of on-beads DNaseI cleavage assay.

## Abbreviations

TEs: Transposable elements
DBD: DNA binding domain
DmTNP: Drosophila P element transposase
THAP: Thanatos Associated Proteins

## Authors’ contributions

HMS initialized the study and generated the data. HMS & SM analyzed the data. SM & HMS wrote the manuscript. All authors read and approved the final manuscript.

## Acknowledgements

We acknowledge Dr. Umashankar Singh for providing suggestions and reviews about the study, Mr. Divyesh Patel and Manthan Patel for help with troubleshooting for western results, Dr. Dhiraj Bhatia Tarushyam Mukherjee and Pravin Hivare for help with image capture and processing using the Leica confocal microscope and ImageJ software.

